# Granulocyte Derived Resistin Inhibits Monocyte Maturation and Induces Immune Suppression in CMML

**DOI:** 10.1101/2025.03.03.640303

**Authors:** Nathan J Hull, Rachel Cant, Laura A Guest, Yu-Hung Wang, Kristian Gurashi, Roberto Paredes, Hector Huerga Encabo, Chien-Chin Lin, Hwei-Fang Tien, Robert Sellers, Sudhakar Sahoo, Dominique Bonnet, Dorothee Selimoglu-Buet, Eric Solary, Daniel H Wiseman, Kiran Batta

## Abstract

Chronic myelomonocytic leukaemia (CMML) is a haematological malignancy characterised by overlapping myeloid dysplasia and proliferation with persisting monocytosis. While monocytes are the cardinal malignant cell type in CMML, as a stem cell neoplasm the disease clone comprises most lineages and differentiation stages, including granulocytes. To investigate the pathogenic contribution of granulocytes in CMML maintenance and progression, we performed phenotypic, transcriptomic and functional characterization of CMML granulocytes. Compared with healthy age-matched controls, CMML granulocytes exhibit defective maturation with reduced granularity and phagocytic capacity. Transcriptome analysis revealed activation of pathways linked to proliferation, Myc activity and inflammation. Notably, *RETN*, which encodes the inflammatory mediator resistin, was upregulated approximately 100-fold in CMML granulocytes; but not differentially expressed in CMML PBMNCs, sorted monocytes, or stem and progenitor cells compared to their healthy counterparts. Accordingly, resistin protein levels were 10-fold higher in plasma from CMML patients and higher plasma resistin levels correlate with poor overall survival and AML-free survival. Remarkably, exposure of healthy monocytes to exogenous recombinant resistin inhibited monocyte maturation and macrophage differentiation. Transcriptome analysis of resistin treated monocytes revealed that resistin induces gene signatures related to immune suppression and myeloid-derived suppressor cell phenotype. We found *SEMA4A* to be a downstream target of resistin and overexpressed in CMML monocytes. Consistent with known roles for SEMA4A, CMML patients displayed higher percentage of Tregs and elevated Th2/Th1 ratio compared with healthy controls and percentage of Tregs corresponding with associated resistin levels. Furthermore, we demonstrated that resistin directly skews the Th2/Th1 ratio via binding to monocytes. In conclusion, we showed that immature granulocytes in CMML produce high levels of resistin, which contributes to defective monocyte maturation and immune suppression.

## Introduction

Chronic myelomonocytic leukaemia (CMML) is a clonal haematopoietic stem and progenitor cell (HSPC) disorder, characterised by features overlapping those of myelodysplastic syndromes (MDS) and myeloproliferative neoplasms (MPN). Persistent monocytosis, with peripheral blood (PB) monocytes ≥0.5 x 10^9^/L and ≥10% of total white blood cells, is a defining feature of CMML^1^.

While monocytes are the driving malignant cell type in CMML, as a stem cell neoplasm the disease clone comprises most lineages and differentiation stages, with morphological dysplasia and functional defects recurrent features beyond the monocyte compartment^2^. Specifically, granulocyte abnormalities are frequent in CMML patients, with neutrophilia, neutropenia, morphological dysplasia and defective microbicidal function common features^3,4^. However, the role these cells play in crosstalk with other immune cell populations, and how they contribute to the disease phenotype, are largely unknown. Under normal development, granulocyte precursors (including promyelocytes, myelocytes and metamyelocytes) develop within the bone marrow (BM) and extravasate when fully matured into the terminal effector cell^5^. An accumulation of granulocyte precursors has been found in PB of CMML patients^6^. Immature granulocytes tend to be proliferative, resistant to spontaneous apoptosis and are often thought to be granulocyte-derived myeloid derived suppressor cell (MDSC) type with immune suppressive properties^7,8^. Immune suppressive cell types are observed in many myeloid leukemias including CMML^9^. Recently we showed that monocytes from CMML patients preferentially polarize towards the immune suppressive M2 macrophage phenotype^10^. Consistent with this, CMML patients have elevated Th2/Th1 ratio, and CD4 terminal effector cell populations compared to healthy individuals^10,11^. It has also been shown that CMML patients with BM dendritic cell aggregates have expanded systemic immune suppressive Treg compartments^11^. Together these reports suggest an immunological bias towards the suppressive state within CMML.

Granulocytes secrete a variety of immunoregulatory factors through which they regulate both innate and adaptive immune systems. In cancer patients, distinct populations of neutrophils at different stages of development are found both in PB and the tumor microenvironment^8^. In solid cancers, tumor-associated neutrophils (TANs) play contrasting roles in tumor progression and can be polarized to N1 and N2 subtypes, analogous M1 and M2 macrophage subtypes, with anti- and pro-tumor functions respectively^12^. The varied secretome profiles of different types of granulocytes (mature vs immature; N1 vs N2) are likely to modulate immune responses differently via crosstalk with other immune cell types. Indeed, in CMML, immature granulocytes secrete alpha-defensins 1-3 (HNP1-3), which can inhibit monocyte differentiation to macrophages^7^. In a recent study, the release of CXCL8 from CMML granulocytes was shown to inhibit the proliferation of wild-type HSPCs, but not mutated HSPCs from CMML patients, suggesting a pathogenic role for CMML granulocytes^13^. However, the impact of CMML granulocytes on immune responses in CMML and the mechanisms involved are unknown.

Here, we investigated the role CMML granulocytes may have in the contribution to disease phenotype and progression. Through flow cytometry, transcriptomic and functional analyses, we present the immature and defective functional status of CMML granulocytes. We found that CMML granulocyte produce elevated levels of cytokine, resistin, which inhibits monocyte maturation and promotes the expansion of immune suppressive cell types by interacting with monocytes via TLR4.

## Results

### CMML granulocytes are enriched for immature forms

To investigate phenotypic and functional alterations of granulocytes in CMML we isolated granulocytes from freshly collected PB from healthy volunteers (n=19) and CMML patients (n=26), using a negative selection method (purity >90%) (Figure 1A, Table S1). Flow cytometry analysis measuring size and granularity of sorted granulocytes from CMML patients displayed significantly decreased granularity compared to healthy controls, suggesting a relatively immature status of CMML granulocytes (Figures 1B & 1C). Consistent with the decreased granularity, the percentage of CD11b/CD16 double positivity, a phenotype marker of mature granulocytes, was significantly lower in CMML patients compared to controls (Figures 1D & 1E). Further, varying percentages of CD11b^+^/CD16^-^ granulocytes were observed in CMML patients, suggesting variability of maturation status in CMML (Figure S1A). Expression levels of CD66b peaks during intermediate stages of granulocyte maturation^14^ and we found significant increase in CD66b expression levels in CMML granulocytes, again consistent with more immature status (Figure 1F). We also observed a marginal but significant increase in expression of CD11b in CMML granulocytes (Figure S1B).

**Figure 1:**
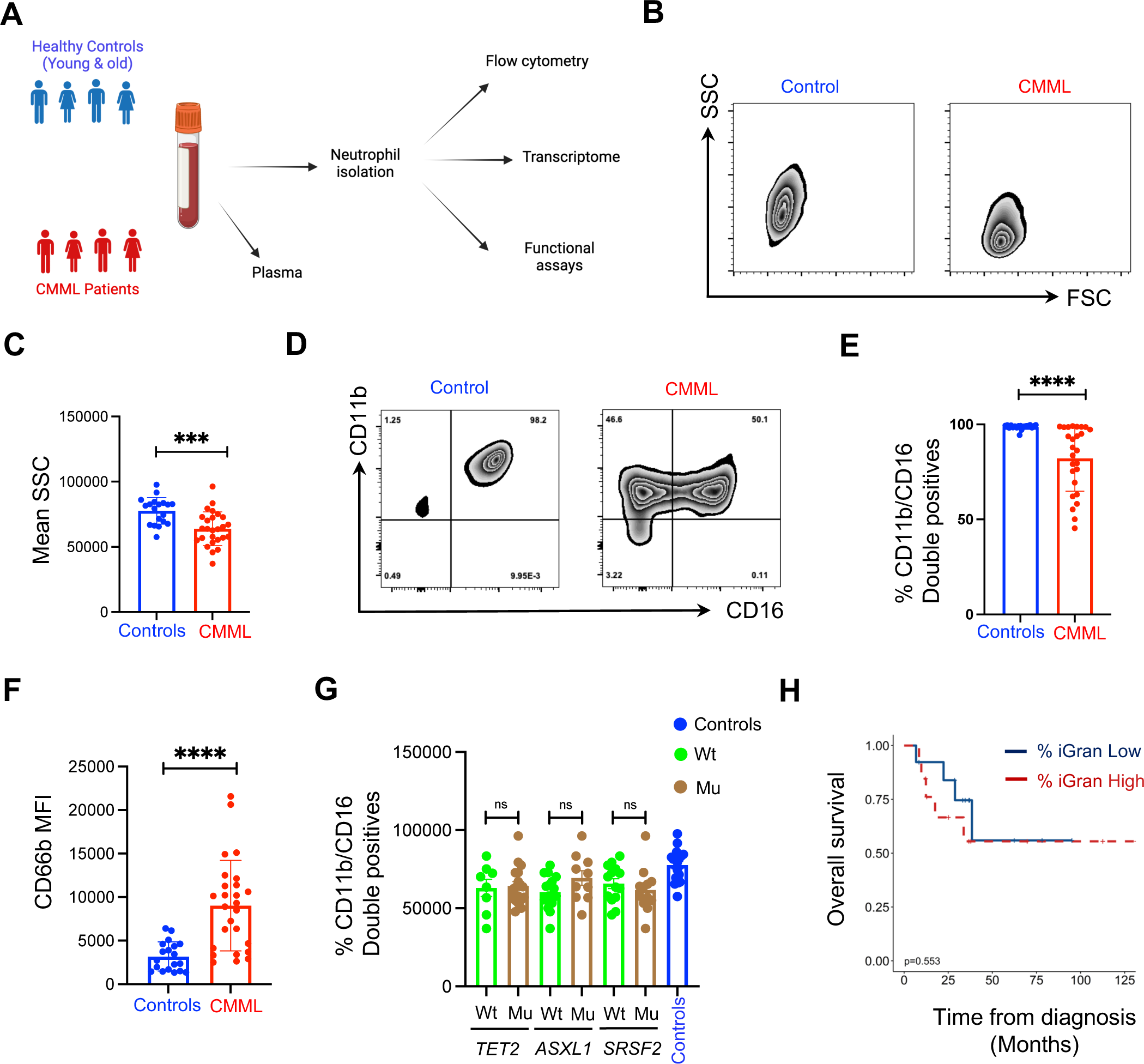
CMML granulocytes are immature: **A.** Schematic diagram outlining the study investigating molecular and functional characterization of CMML and healthy control granulocytes. **B.** Representative flow cytometry plots displaying the granularity and size of control and CMML granulocytes. **C.** Bar graph showing the mean SSC between control (N=19) and CMML granulocytes (N=26). **D.** Representative flow cytometry plots displaying the expression of CD11b and CD16 on control and CMML granulocytes. **E.** Bar graph showing average percentage of CD11b/CD16 double positive cells in control (N=19) and CMML (N=26) granulocytes. **F.** Average mean fluorescence intensity (MFI) of granulocyte marker CD66b in control (N=19) and CMML (N=26) granulocytes. **G.** Bar graph showing percentage of CD11b/CD16 double positive mature granulocytes in wild-type and mutant genotype of indicated genes recurrently observed in CMML patients. **H.** Kaplan-Meier overall survival plot of CMML patients stratified by the percent of immature granulocytes (iGran) classified based on CD11b^+^CD16^-^ phenotype. High = > 13% (N=13) vs low = <13% (N=13). Error bars = ± SEM; ns, no significance; ***, P<0.001; ****, P<0.0001.

Given the observed heterogeneity in maturation status of CMML granulocytes and previous reports linking loss of *TET2* and *ASXL1* to granulocyte maturation defects, we explored if CMML granulocyte phenotype is linked to mutation status^15,16^. Our results showed no significant difference in percentage of mature granulocytes or mean SSC between mutant and wild type CMML granulocytes for the genes most recurrently mutated in CMML (Figures 1G & S1C). This suggests that decrease in granularity and the presence of immature granulocytes are associated with all three frequently occurring mutations in CMML. Additionally, we did not find significant differences in maturation status or granularity between CMML patients with dysplastic versus proliferative features (Figure S1D). Next, we examined whether presence of immature granulocytes is linked to prognosis but found no differences in overall survival or AML-free survival between patients with high percentage of CD11b^+^/CD16^-^ immature granulocytes (>13%) vs low percentage of CD11b^+^/CD16^-^ immature granulocytes (<13%) (Figure 1H & S1E). Having not observed any specific associations with genotype, we next explored whether low density granulocytes (LDGs) carry higher mutation burden over high density granulocytes (HDGs) within the same patient. We sorted HDGs and LDGs from four CMML patients and sequenced for most commonly occurring mutations in myeloid cancers (Figure S1F). We found no differences in variant allele frequencies for known mutations between HDGs vs LDGs for any of the four patients (Figure S1G). Overall, we found CMML granulocytes to be less granular and more immature compared with healthy controls, an observation independent of mutation status, CMML type or disease prognosis.

### CMML granulocytes are transcriptional distinct from their healthy counterparts

To define gene expression programme changes in CMML granulocytes we performed RNA-seq on isolated CMML (n=14) and healthy (n=7 young, n=7 aged) granulocytes. Principle component analysis (PCA) clearly segregated CMML granulocytes from healthy controls, but not healthy granulocytes based on age (Figure 2A). When analysing aged and young granulocytes separately, we observed 378 genes (180 up and 198 down) to be differentially expressed (Table S2). Upregulated genes in aged granulocytes, in comparison with young granulocytes, include negative regulator of neutrophil activation *CLEC12A*^17^, UDP-Glucose receptor *P2RY14* that promotes inflammation in neutrophils^18^ and transcription factor *IRF1* whose expression is linked to neutrophil maturation^19^. The key regulator of neutrophil trafficking *CXCR4*^20^, ER-lipid raft associated protein *ERLIN2* and neutrophil motility regulator *NPC1*^21^ were all downregulated in aged neutrophils (Figure S2A). Comparing CMML granulocytes with granulocytes from healthy controls (aged and young together) we observed 1375 genes to be differentially expressed (1093 up and 282 down) (Table S3). Pathway analysis of upregulated genes revealed activation of cell cycle, Myc targets and oxidative phosphorylation, among others (Figures 2B & 2C). Downregulated genes are involved in inflammation, apoptosis and interferon responses pathways (Figure S2B). Indeed, proapoptotic genes such as *BID, FAS* and *TNFSF10* were downregulated in some patients, suggesting active proliferative status of CMML granulocytes (Figure S2C).

**Figure 2:**
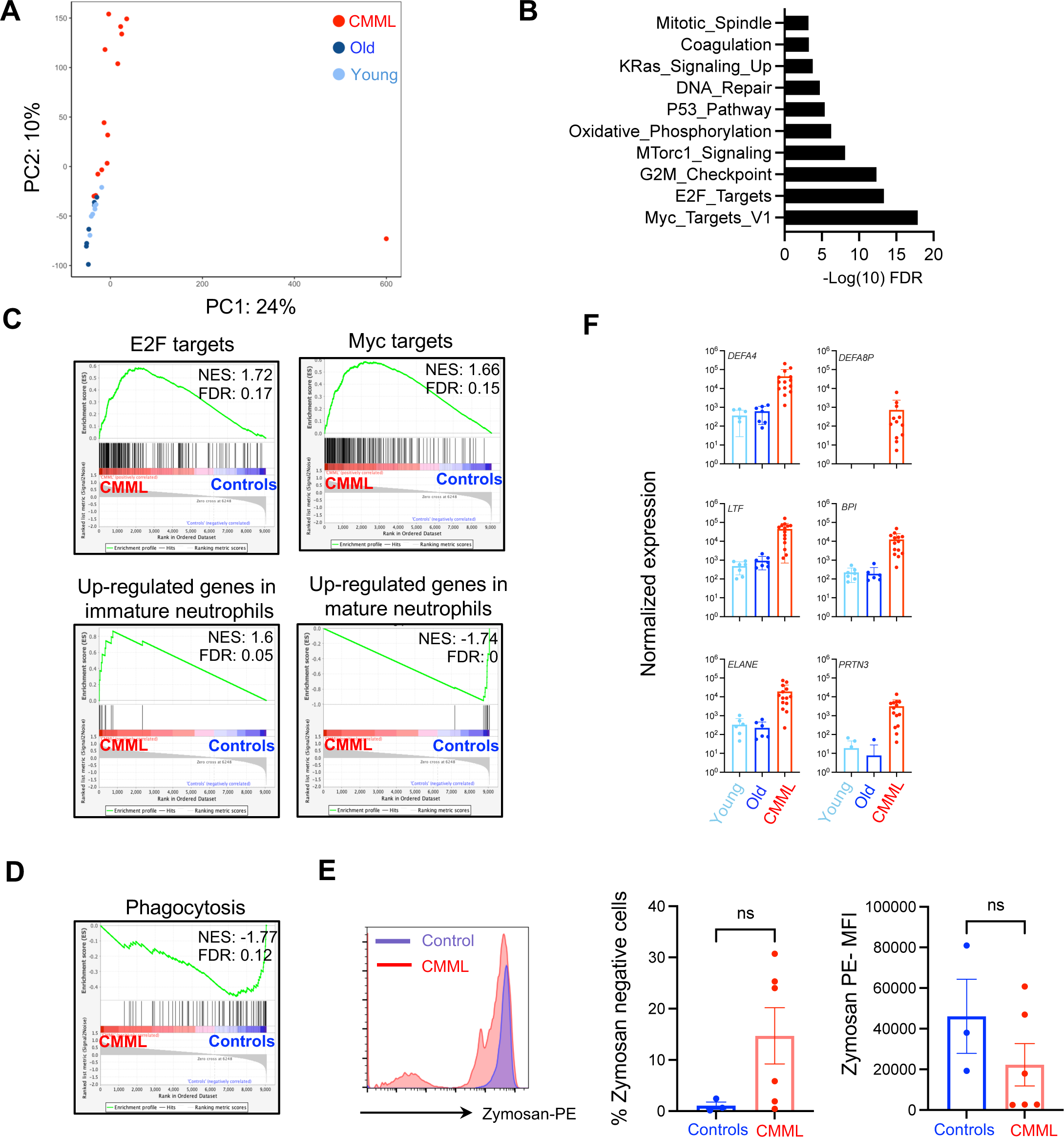
Transcriptional profile of CMML granulocytes: **A.** Principal component analysis of RNA-seq data performed on healthy young (N=7) and old control (N=7) and CMML (N=14) granulocytes. **B.** Gene ontology analysis of upregulated genes in CMML granulocytes as compared with all controls. Top ten significant pathways are shown. **C.** GSEA plots showing relative enrichment of indicated gene sets in CMML granulocytes as compared with controls. **D.** GSEA plot showing relative enrichment of gene set related to phagocytosis in CMML granulocytes as compared with controls. **E.** A representative flow cytometry plot displaying the phagocytic uptake of zymosan particles by control (N=3) and CMML (N=6) granulocytes (left). Bar graphs depicting the percentage of zymosan negative cells (middle) and mean fluorescence intensity of zymosan particles (right) in phagocytosis uptake assays performed with healthy and CMML granulocytes. **F.** Normalized expression levels of indicated genes in healthy young and old and CMML granulocytes. Error bars = ± SEM; ns, no significance.

To validate the immature status of CMML granulocytes seen by flow cytometry, we performed Gene Set Enrichment Analysis (GSEA) using gene sets differentially expressed between healthy mature and immature neutrophils^22^. Genes upregulated in immature neutrophils were positively regulated in CMML granulocytes and, conversely, genes upregulated in mature neutrophils were negatively regulated (Figure 2C), supporting immature status of CMML granulocytes. Given that granularity and maturation status of neutrophils are linked to key neutrophil functions (e.g. phagocytosis), we investigated phagocytic capacity of CMML granulocytes compared with controls. GSEA also suggested that genes involved in phagocytosis are negatively regulated in CMML granulocytes compared with controls (Figure 2D). To validate this, we performed a phagocytic uptake assay using zymosan-PE beads in healthy (n=3) and CMML granulocytes (n=6). We found a near 100% positive uptake in healthy controls, however, varying fractions of CMML granulocytes found to be defective in phagocytic capacity (Figure 2E).

The top 20 significantly upregulated genes included defensins *(DEFA4, DEFA8P)*, genes involved in neutrophil degranulation and NETosis *(ELNAE, PRTN3, MPO)* and genes associated with maturation status *(LTF, BPI)* (Figure 2F). Top downregulated genes include *CXCR2* and *CXCR1*, both involved in neutrophil activation and recruitment, and the G protein coupled receptor *KISS1R* (Figure S2D). Consequently, signalling by CSF3, a pathway involved in neutrophil maturation and production, is negatively regulated in CMML granulocytes in comparison with controls (Figure S2E). Other genes downregulated and linked to neutrophil activation and antimicrobial activity included *SOD2*, *CSF3R* and *MNDA* (Figure S2F). Taken together these results confirm the immature status of CMML granulocytes observed phenotypically and identify several functionally important pathways defective in CMML granulocytes, including phagocytosis.

### Resistin is upregulated specifically in CMML granulocytes

One of the top upregulated genes but with little known role in neutrophil function was *RETN*, which encodes the adipokine resistin. First, we asked whether *RETN* upregulation is unique to neutrophils in CMML, as compared with other lineage populations forming the disease clone. Interrogating available transcriptome data^23,24^, we found *RETN* is uniquely upregulated in CMML granulocytes but not in disease initiating stem cells, disease propagating monocytes or peripheral blood mononuclear cells (PBMNCs) (Figure 3A). Next, we investigated whether upregulation of *RETN* in CMML granulocytes is unique to CMML, as compared with other myeloid cancer types. We thus re-analysed publicly available BEAT-Neutrophil datasets^25^ and found *RETN* levels are also elevated in neutrophils from chronic myeloid leukaemia (CML), chronic neutrophilic leukaemia (CNL), atypical CML, MDS/MPN-unclassifiable and MPN-unclassifiable patients (Figure 3B).

**Figure 3:**
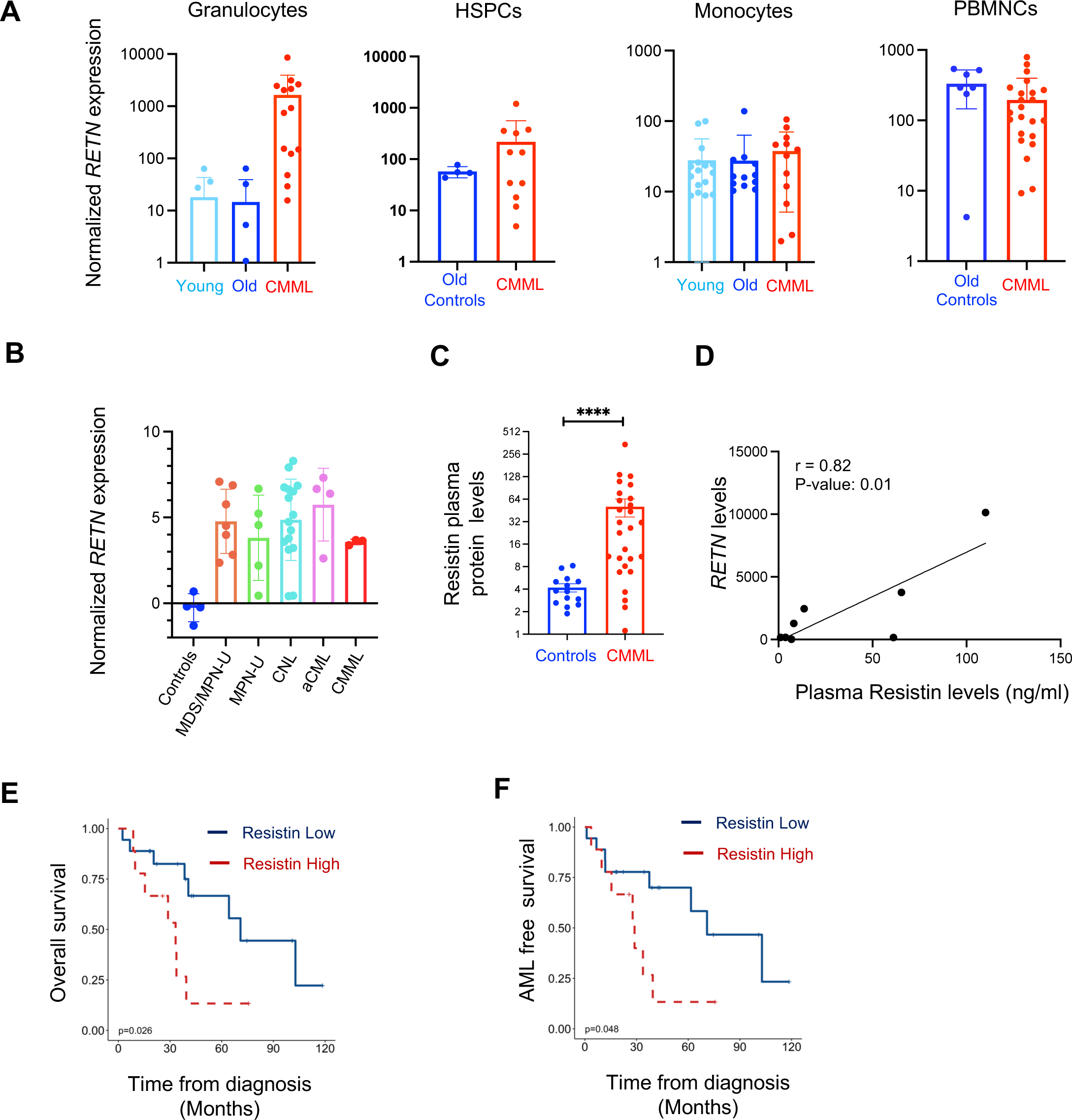
Resistin is upregulated specifically in CMML granulocytes: **A:** Normalised expression levels of *RETN* in granulocytes, CD34+ HSPCs, CD14+ monocytes and whole PBMNCs from CMML patients and healthy (young and/or old) controls^23,24,69^. **B.** Relative expression levels of *RETN* in granulocytes sorted from different myeloid cancer patients^25^. aCML: atypical CML, MPN-U: MPN-unclassifiable, CNL: Chronic Neutrophilic Leukaemia, MDS/MPN-U: MDS/MPN-unclassifiable. **C.** Resistin protein (ng/ml) levels in healthy (N=14) and CMML (N=27) plasma samples as determined by ELISA. **D.** Pearson correlation between CMML granulocyte *RETN* expression and matched plasma resistin concentration. **E & F.** Kaplan-Meier overall (E) and AML-free survival plot (F) of CMML patients stratified by plasma resistin levels (High resistin N=9; Low resistin N=18). Error bars = ± SEM; ns, no significance; ****, P<0.0001.

*RETN* is minimally expressed in stem cells and its expression gradually increases during granulomonocytic commitment, finally peaking at the monocyte stage (Figure S3A, data analysed from Bloodspot). To confirm *RETN* upregulation at the protein level we measured resistin plasma concentration comparing aged healthy volunteers (n=14) and CMML patients (n=27), revealing mean 12-fold higher resistin levels in CMML plasma (Figure 3C). We also found a significant positive Pearson correlation (r=0.82) between *RETN* RNA expression in granulocytes and plasma resistin protein concentration in CMML, suggesting that granulocytes are the major source of elevated resistin levels in CMML plasma (Figure 3D). Interestingly plasma resistin levels are also elevated in individuals experiencing acute bacterial infections, suggesting an important role for resistin in inflammation (Figure S3B)^26^.

Since we observed low-density immature granulocytes in CMML, we sought to determine whether plasma resistin levels were associated with granularity or maturation status. While granularity was not correlated with resistin levels, we found an inverse Pearson correlation between maturation status, as measured by CD11b and CD16 expression, and resistin levels (Figures S3C & S3D). This suggests that immature granulocytes may be the primary source of elevated plasma resistin levels. Next, we investigated whether resistin levels are correlated with any clinical parameters. Whilst plasma resistin levels were not linked to genotype (Figure S3E), interestingly we found that CMML patients with high plasma resistin (n=9) had worse overall survival (OS) compared to those with low plasma resistin (n=18), with median OS 33.2 months and 70.9 months, respectively (Figure 3E). This correlation was extended further with high plasma resistin CMML patients having a worse median AML-free survival of 28.8 months compared to 70.9 months for low plasma resistin patients (Figure 3F). Taken together, these results imply that the high resistin levels found in CMML plasma derives from immature granulocytes, and elevated resistin levels might contribute towards CMML progression.

### Resistin maintains monocyte state via its binding to TLR4

Ex vivo culture of monocytes induces maturation of CD14^+^/CD16^-^ cells to CD14^+^/CD16^+^ state ^27,28^. To assess whether resistin plays any role in monocyte maturation that may contribute towards the CMML phenotype we treated aged healthy monocytes, ex vivo, with increasing concentrations of resistin or GFP endotoxin control for 24 and 48 hours. Surprisingly, resistin prevented transition of CD14^+^/CD16^-^ monocytes to CD14^+^/CD16^+^ state in a dose dependent manner at 24 hours (Figures 4A & 4B). Even after 48 hours, while most control GFP treated cells progressed to CD14^+^/CD16^+^ state, 250ng/ml resistin treated cells predominantly maintained a CD14^+^/CD16^-^ monocyte state (Figures S4A & S4B). Human resistin has been shown to bind to TLR4 on monocytes and plays a role in immune modulation^29,30^. To investigate whether resistin’s binding to the TLR4 receptor is important for the observed phenotype, aged healthy monocytes were treated with TLR4 neutralising antibody (TLR4Ab) prior to resistin treatment. We found resistin’s ability to maintain CD14^+^/CD16^-^ monocyte state was blocked by the addition of TLR4Ab (Figures 4C & 4D). Again, this effect was also observed after 48 hours treatment, with ∼66% CD14^+^/CD16^-^ cells in resistin treated cells compared to ∼13% with addition of TLR4 nAb and ∼15% in control GFP treated cells (Figures S4C & S4D). Coincidently, analysis of publicly available datasets revealed higher expression levels of *RETN* in CD14^+^/CD16^-^ monocytes compared to other monocyte subtypes suggesting a possible cell autonomous role for resistin is maintaining CD14^+^/CD16^-^ monocyte state (Figure 4E)^31^.

**Figure 4:**
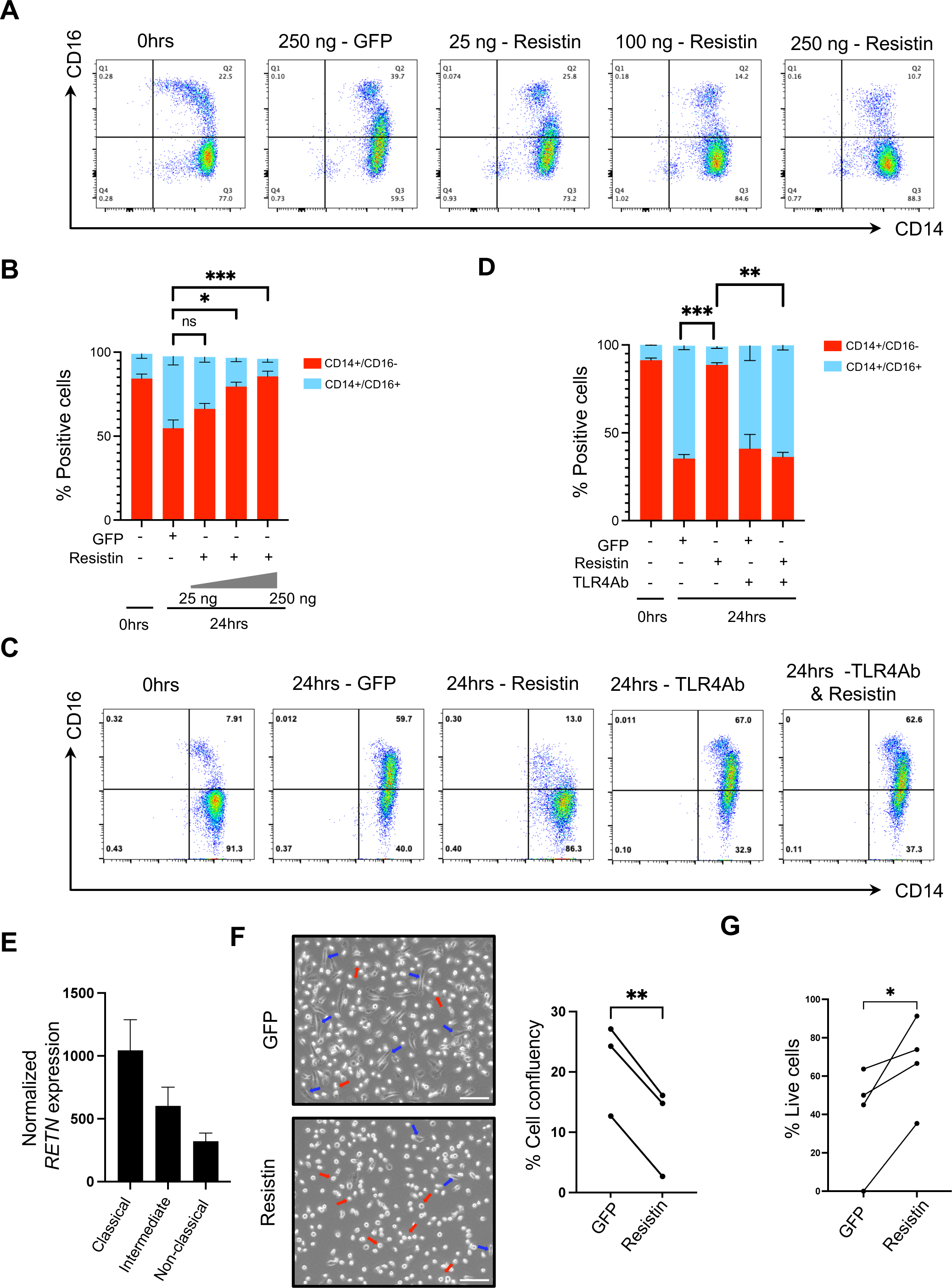
Resistin inhibits monocyte maturation. **A.** Representative flow cytometry plots showing the surface marker expression of CD14 and CD16 on freshly isolated healthy monocytes (0hrs) and monocytes treated with increase in concentration of resistin or control GFP for 24 hours (N=5). **B & D.** Bar charts showing average percentage of CD14+ or CD14+CD16+ populations under indicated conditions. **C.** Representative flow cytometry plots showing the surface marker expression of CD14 and CD16 on monocytes incubated with or without TLR4 neutralising antibody (TLR4Ab) prior to exposure with resistin or GFP control for 24 hours (N=4). **E.** Relative expression levels of *RETN* in three different types of human healthy monocytes^31^. **F.** Representative brightfield microscope images of healthy monocytes/macrophages morphology following 6-day differentiation of monocytes treated either with 250 ng/ml of GFP control or resistin (Left). Blue arrows = attached macrophages; red arrows = suspended monocytes; scale bar 100 µm. Quantification cell confluency after 6 days treatment of monocytes either with GFP control or resistin (right). **G.** Percentage of live cells in suspension cultures of monocyte differentiation, treated either with GFP control or resistin. Error bars = ± SEM; ns, no significance; *, P<0.05; **, P<0.01; ***, P<0.001.

CMML monocytes are resistant to apoptosis by upregulation of MCL1^32^. We asked whether resistin promotes the survival of monocytes and thereby contributes to the defining monocytosis observed in CMML. To this end, we measured apoptosis levels by Annexin V staining following treatment with either resistin or GFP control in CMML monocytes but saw no changes in the percentage of apoptotic cells following resistin treatment (Figures S4E & S4F). We also measured cell growth following treatment with resistin and again we found that resistin does not promote CMML monocyte survival (Figure S4G). Together these results suggest that while resistin maintains monocyte state, it does not directly contribute to the absolute monocytosis.

Defects in activation and maturation to macrophages have been reported in CMML monocytes^6,33^. To investigate whether resistin affects monocyte differentiation to macrophages we cultured healthy monocytes, *ex vivo*, in presence of resistin and M-CSF for 6 days. We observed that monocytes failed to attach and differentiate into macrophages in the presence of resistin, while control GFP-treated monocytes robustly differentiated into macrophages (Figure 4F). To rule out the possibility that resistin might be killing monocytes and thereby preventing them to attach and differentiate, we quantified percentage of live cells that were in suspension and free-floating. Our results showed more live cells in the resistin-treated condition compared to GFP control, suggesting again that resistin does not impact monocyte survival (Figure 4G). Together, these findings show that resistin inhibits monocyte differentiation via the TLR4 signalling pathway and maintains them in CD14^+^/CD16^-^ monocyte state.

### Resistin induces transcriptional alterations in inflammatory pathways

To investigate direct transcriptional changes induced by resistin we treated healthy monocytes (N=4) with either recombinant resistin protein or control GFP protein for 6 and 24 hours. Freshly isolated healthy monocytes, before culturing, were also used as controls (0 hrs). Since resistin and LPS both bind to TLR4, we aimed to investigate transcriptional changes specifically induced by resistin by using LPS treated monocytes as an additional control (Figure 5A). PCA analysis of transcriptome data clearly segregated treated samples from 0 hrs controls. Resistin-treated samples were placed between GFP control and LPS treated samples suggesting distinct transcriptional changes induced by resistin (Figure 5B). We found 923 (528 up, 396 down) and 773 (483 up, 189 down) genes differentially expressed at 6 and 24 hrs, respectively, in resistin treated cells as compared to GFP controls (Tables S4, S5). Total 222 genes were upregulated at both time points, including *DEFA1, PDGFRB1, CXCL1, IL1* and *CCL5*. Notably alfa-defensin (HNP1-3) encoding genes (e.g. *DEFA*), which are known to inhibit monocyte maturation, were also upregulated in resistin treated monocytes (Figure S5A)^6,13^. The number of genes downregulated at 24 hours was significantly reduced from 6 hrs (396 vs 189) and only 21 genes were commonly downregulated at both time points (Figure S5B). These include the myeloid cell nuclear differentiation antigen *MNDA*, epigenetic factor *IDH2*, & hepatocyte growth factor *HGF1,* all of which positively regulate monocyte activation and macrophage differentiation^34,35^. Analysis of upregulated genes upon resistin treatment revealed activation of inflammatory responses, cytokine production and cell motility (Figures 5C & S5C). GSEA analysis further confirmed activation of TNF-A signalling in resistin treated cells compared to controls (Figure 5D).

**Figure 5:**
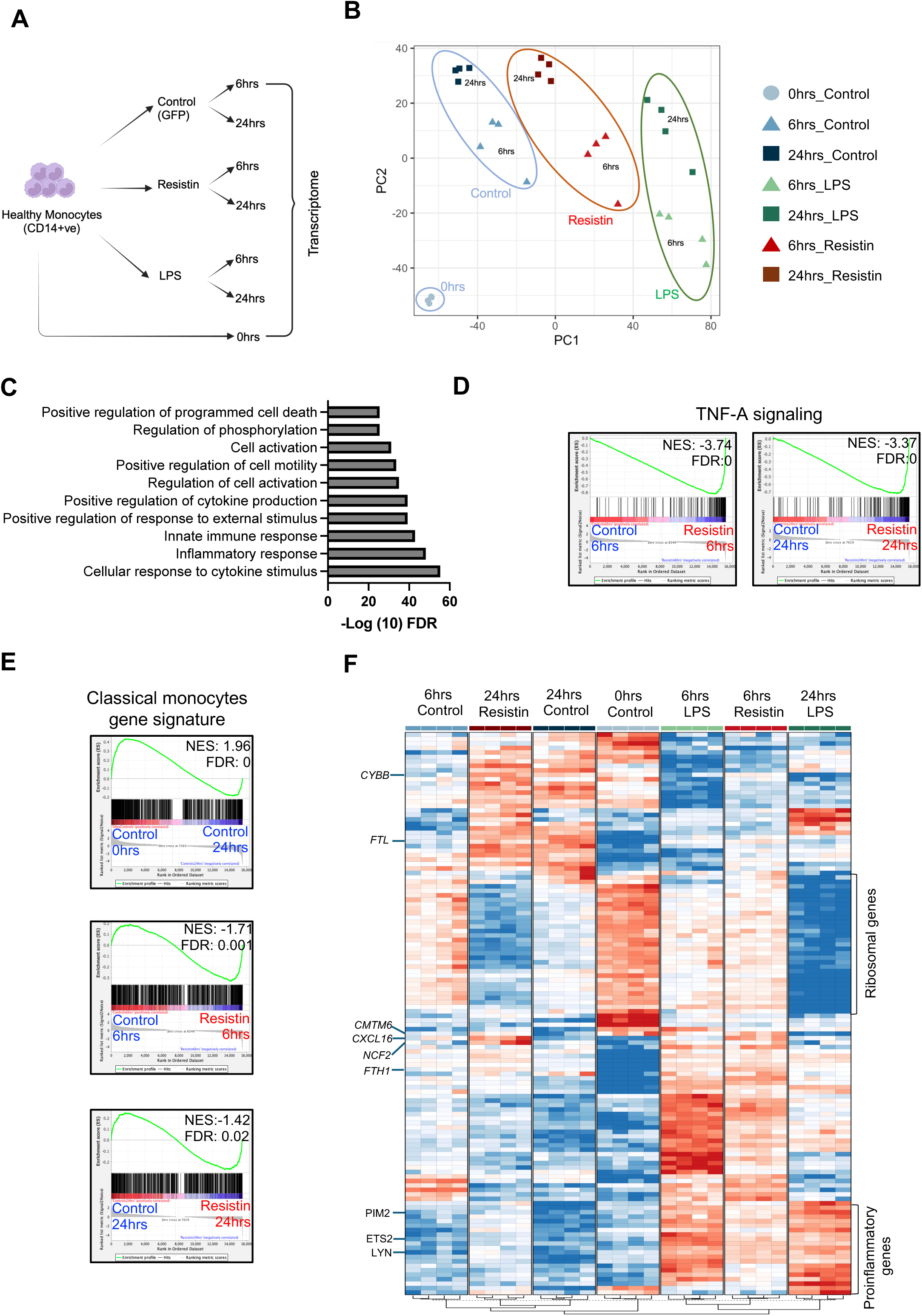
Resistin maintains monocyte gene expression signature: **A.** Schematic diagram depicting transcriptome study investigating downstream regulators of resistin or LPS on healthy monocytes. Monocytes treated recombinant protein GFP protein and freshly isolated monocytes were used as controls. **B.** Principal component analysis of the RNA-seq data. **C.** GO analysis of genes induced by resistin at 6 and 24 hrs together as compared with respective time point controls. Top ten significant pathways are shown. **D & E.** GSEA plots showing relative enrichment of indicated gene sets in different comparisons as listed. **F.** Heatmap showing expression of top30 DEGs across different comparisons.

Cell surface marker analysis of resistin treated cells suggested that resistin maintains cells in monocyte state (Figure 4). We next wanted to confirm whether resistin inhibits monocyte maturation at gene expression level. To this end we compared the expression levels of genes known to be uniquely expressed in classical monocytes^31^, which represents the majority of healthy monocytes, in resistin treated cells. The classical monocyte gene set was negatively regulated as cells start to differentiate i.e. 0 hrs vs 24 hours GFP treated (Figure 5E, top panel). However, resistin treated cells maintained this gene expression programme at both 6 hrs and 24 hrs as compared to their respective timepoint controls, further supporting resistin’s role in maintaining monocyte state (Figure 5E, bottom two panels).

Next, we investigated the top 30 DEGs across different comparisons, to identify dynamic gene expression patterns between samples (Figure 5F). We found proinflammatory genes to be upregulated in both LPS and resistin treated cells, albeit to a lesser extent in resistin treated cells (Figure 5F). Conversely, ribosomal genes were downregulated in both LPS and resistin treated cells, again more profoundly in LPS treated cells. Many genes linked to inflammation and monocyte adhesion, including *ETS2, LYN, & PIM2*, were transiently upregulated by resistin but consistently by LPS^36–38^. In contrast, *CXCL16, NCF2, CYBB* levels were higher in resistin treated monocytes as compared to LPS; these genes are involved in autoimmunity, atherosclerosis & foam cell maturation, consistent with resistin’s established role as a biomarker of atherosclerosis^39–41^. Ferritin subunits *FTL* and *FTH1* were also induced by resistin, and their expression is linked to immune suppression in many cancers^42^. Taken together these results suggest that resistin exposure predominantly upregulates inflammatory gene expression, maintains monocyte gene expression program and downmodulates monocyte maturation gene signatures.

### Resistin induces immune suppression in CMML

To investigate genes uniquely induced by resistin, we overlapped upregulated genes that were induced by either resistin or LPS at both 6 and 24 hours.

This identified a total of 99 genes uniquely regulated by Resistin at either time point, of which 6 were upregulated at both time points. These included *SEMA4A, CCL24, TMEM138, MS4A7, EFHD2* and *CMTM6* (Figure 6A, Figure S6A). Remarkably, all these genes have known roles in inducing immune suppression and M2 macrophage polarization^43–46^. Next, we overlapped genes uniquely induced by resistin with those upregulated in CMML monocytes^23^ reasoning that common genes might be relevant targets of resistin in CMML. We found 12 genes common between these datasets, including *SEMA4A, SDC2, TMEM138,* and *SPP1* (Figure 6B). Of note, *SEMA4A* is upregulated at both 6 and 24 hrs following treatment with resistin and is also upregulated in CMML monocytes (Figures 6C and 6D). We further validated that SEMA4A expression is indeed induced at the protein level following treatment with resistin, but not by control GFP protein (Figure 6E). Furthermore, *SEMA4A* and *RETN* levels are positively correlated in AML samples, suggesting that resistin could be an upstream mediator of *SEMA4A* expression (Figure S6B)^47^.

**Figure 6:**
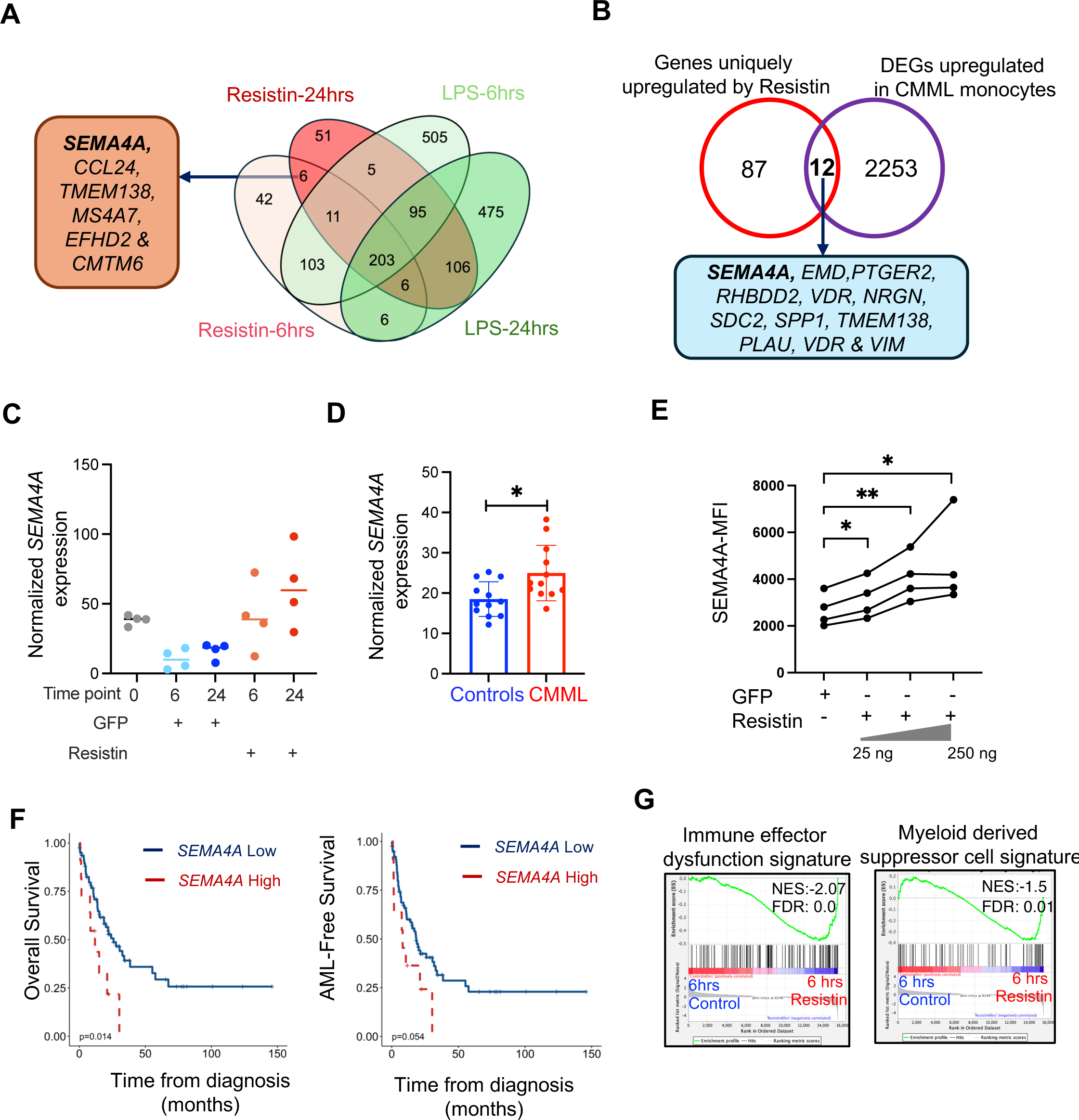
Resistin unique targets in CMML monocytes are linked to immune suppression: **A.** Venn diagram showing overlap of genes upregulated following 6 and 24 hrs treatment with either resistin or LPS in comparison with their respective time point controls. **B.** Venn diagram showing overlap of genes uniquely upregulated by resistin but not by LPS with genes that are upregulated in CMML monocytes^23^. **C.** Relative expression levels of *SEMA4A* in monocytes treated with either resistin or control GFP protein. **D.** Relative expression levels of *SEMA4A* in CMML and healthy monocytes. **E.** Mean fluorescence intensity (MFI) of SEMA4A on healthy monocytes treated with either control GFP or increase in concentration of resistin for 24 hrs (N=4). **F.** Kaplan-Meier overall (left) and AML-free (right) survival plot of CMML patients stratified by *SEMA4A* expression levels (N=11 High; N=79 Low) **G.** GSEA plots showing relative enrichment of indicated gene sets in resistin (6 hrs) treated monocytes in comparison with the controls. Error bars = ± SEM; ns, no significance; *, P<0.05; **, P<0.01.

In CMML patients, higher *SEMA4A* expression is linked to poor overall survival and a trend towards inferior AML-free survival (Figure 6F). Deconvoluting cell signatures between *SEMA4A*-high and *SEMA4A*-low CMML patient samples suggested that *SEMA4A*-high levels are linked to a decrease in stemness signature and an increase in monocyte and M2 macrophage signatures (Figure S6C). The induction of several genes involved in immune suppression upon resistin treatment, in conjunction with the M2 macrophage signature in *SEMA4A*-high CMML BM-MNCs, suggests defects in immune responses in a subset of CMML patients. Furthermore, GSEA analysis suggested that resistin treatment induces myeloid derived suppressor cell and immune effector dysfunction signatures (Figure 6G). Overall, these data suggest that resistin induces an immune suppression gene signature, presumably via SEMA4A.

SEMA4A has been shown to drive Th2 responses and stabilise the Treg cell phenotype^48,49^. To investigate whether resistin-mediated *SEMA4A* induction alters T cell profiles, we performed *ex vivo* culture of PBMNCs in presence or absence of resistin for 9 days, analysing the resultant Th2/Th1 cell ratio and Treg populations by flow cytometry. As controls, we depleted monocytes from PBMNCs to interrogate whether any changes in T cell profiles are indeed mediated by resistin’s interaction with monocytes. Results showed that resistin elevated the ratio of Th2/Th1 compared to control, only in the presence of monocytes but not when monocytes were depleted (Figure 7A). However, we did not observe any specific expansion of the Treg population in presence of resistin (Figure S7A). Next, to investigate whether resistin plasma levels in CMML patients correlate with either of the T-cell phenotypes we observed in *ex vivo* cultures, we measured the Th2/Th1 ratio and Treg fraction in CMML patients versus healthy controls. Consistent with our previous findings, we observed elevated Th2/Th1 ratio in PB of CMML patients (n=8) compared with healthy controls (n=10) (Figure 7B) ^10,11^. However, CMML patients with high or low levels of resistin both displayed significantly higher Th2/Th1 ratio than the controls (Figure 7C). We also found that CMML patients’ (n=13) PB had a 3-fold higher percentage of Treg cells than healthy controls (n=8) (Figure 7D). Interestingly, CMML patients with higher plasma resistin levels had a significant 4-fold higher Treg percentage than controls (Figure 7E), however, CMML patients with lower plasma resistin levels showed no significant difference in percentage of Tregs compared to healthy controls (Figure 7E). In addition, we found a strong correlation (r=0.83) between plasma resistin concentration and percentage of Tregs in CMML PB (Figure S7B). Taken together these results show that immune suppression is observed in CMML and resistin is a candidate driver of this phenotype.

**Figure 7:**
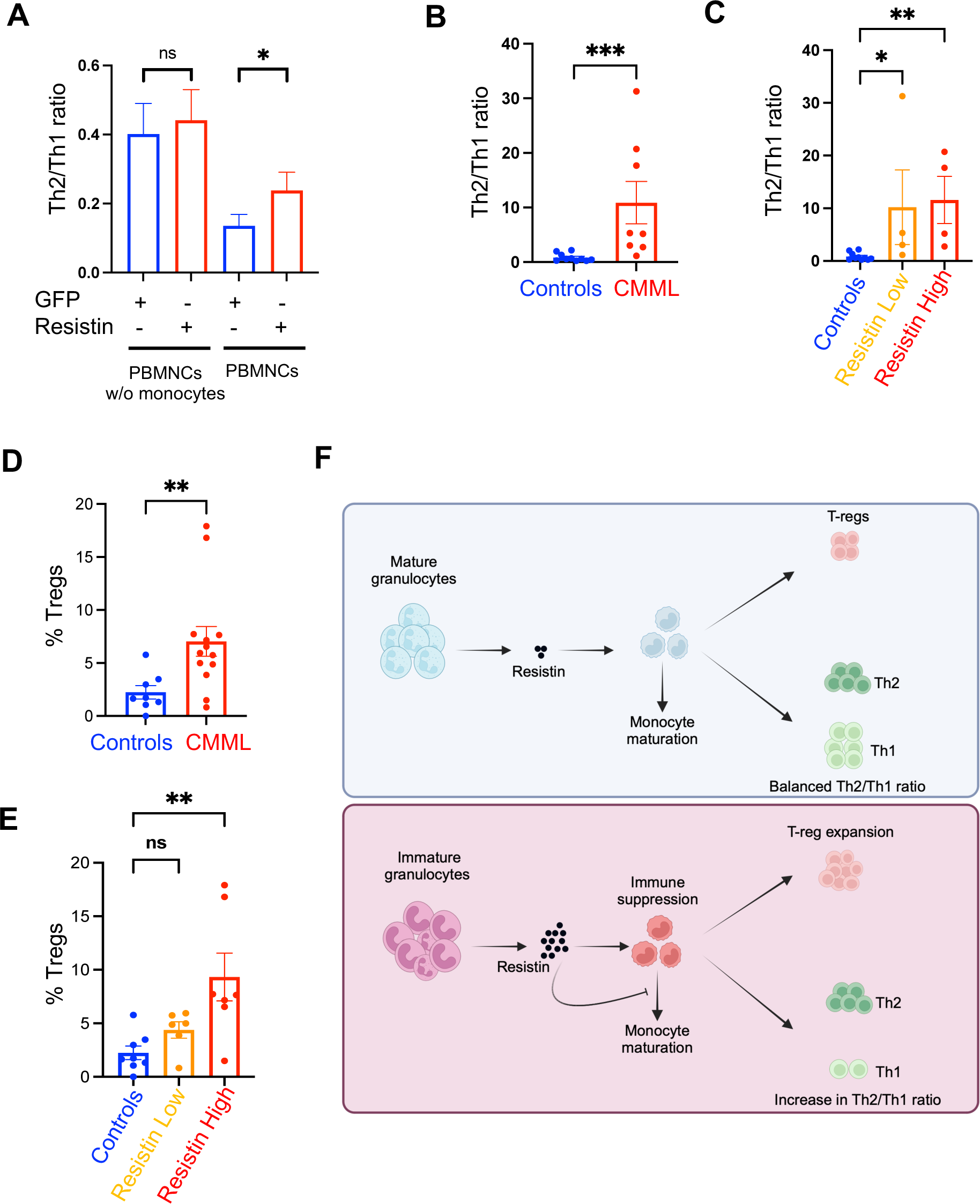
Resistin induces immune suppression in CMML: **A.** Ratio of Th2/Th1 cells in ex vivo cultures of whole- or CD14 depleted-PBMNCs treated with either GFP or resistin for 9 days (N=6). **B.** Th2/Th1 ratio from healthy (N=10) and CMML (N=8) peripheral blood (PB). **C.** Ratio of Th2/Th1 cells in PB of CMML patients stratified based plasma resistin low (N=4) and high levels (N=4). **D.** Percentage of Treg cells in healthy (N=8) vs CMML PB (N=13). **E.** Percentage of Treg cells in PB of CMML patients stratified based on resistin plasma concentration low (N=6) and high (N=7) against healthy controls. **F.** Schematic diagram of the normal monocytic maturation pathway and immune regulation versus dysregulated monocyte maturation and immune suppression found in CMML, caused by resistin that is released by immature granulocytes. Error bars = ± SEM; ns, no significance; **, P<0.01.

In conclusion, we propose a model whereby in healthy individuals mature granulocytes circulate, releasing low basal levels of resistin. CMML patients have increased numbers of immature granulocytes in circulation, which release high levels of resistin into plasma, which inhibits normal monocyte differentiation via binding to TLR4, maintaining them as monocytes and contributing to the monocytosis phenotype in CMML. Resistin further induces an immune suppressive signature in monocytes, leading to increased frequency of Treg cells and an elevated Th2/Th1 ratio, apparently mediated via SEMA4A (Figure 7F).

## Discussion

In this study we show that immature granulocytes contribute to CMML pathophysiology through multiple mechanisms. In addition to being a source of altered chemokines and cytokine profiles, immature granulocytes regulate monocyte activation and differentiation, in part via resistin. Through crosstalk with monocytes, immature granulocytes affect adaptive immune cell profiles and thereby potentially regulate antitumour immunity.

We identified variable fractions of immature granulocytes in CMML patients, in keeping with earlier reports that observed various abnormalities in CMML granulocytes^3,4^. More recently, Deschamps et al. using spectral flow cytometry identified CD15^+^CD16^-^CD66b^+^ immature granulocytes in CMML patients and linked the presence of these cells with poor overall survival and AML-free survival^13^. In our study, we did not observe any association with frequency of immature granulocytes and overall survival. This discrepancy in findings could be due to smaller sample size in our cohort and/or differences in sample preparation. We used negative selection of neutrophils from freshly collected PB to sort for both mature and immature granulocytes, allowing us to investigate all granulocyte populations. By contrast, Deschamps et al. analysed immature granulocytes from PBMNCs isolated following low density gradient centrifugation that separates PBMNCs from mature granulocytes. Similar to their findings, however, we also observed a granulocyte-MDSC signature in CMML neutrophils as measured by the expression of *ARG1*, *MPO*, *S100A8*, *ANXA1* and *S100A12*^50^ and elevated *CXCL8* levels. We also confirmed CMML granulocytes are clonal, carrying all the mutations observed in PBMNCs.

Neutrophils from CMML samples, regardless of the mutation profile, showed reduced granule complexity. This is surprising because murine and human models of *ASXL1* and *TET2KO* in HSPCs resulted in accumulation of higher proportion of immature granulocytes^15,16^. Deschamps et al. showed that mutations in *SRSF2 and ASXL1,* are also linked to this phenotype^13^. Mutations in *TET2, SRSF2* and *ASXL1* together account for more than ∼90% of CMML patients and perhaps this is the reason we did not observe any genotype-phenotype association. Because we observed considerable heterogeneity in maturation status, we reasoned that mature granulocytes were those that did not carry mutations. However, our genotyping results in LDGs vs HDGs argues against this line of thought. Together, these observations suggest that any block in maturation of CMML granulocytes is leaky, mediated by both cell-extrinsic and cell-intrinsic factors. Alternatively, mature neutrophils might be reprogrammed to immature/MDSC state via cell extrinsic factors, for example TGFβ.

The relative abundance of monocyte subsets is altered in various conditions such as infections, cardiovascular disease and cancer^51^. Although classical monocytosis has been established as a key diagnostic feature of CMML, the pathological significance have remained unclear^33^. In this study, we showed resistin keeps monocytes in CD14^+^CD16^-^ state ex vivo via it’s binding to TLR4. However, whether resistin does induce classical monocyte repartition in CMML patients is unresolved and should be investigated. Lack of experimental culture conditions mimicking classical monocyte transition to non-classical monocytes and a functional homology between human and murine resistin precludes testing this directly in ex vivo cultures and in vivo models, respectively. *RETN* levels are also elevated in granulocytes from several myeloid cancer patients, whether monocytes exhibit defects in maturation in these myeloid cancer remains future area of investigation.

Myeloid differentiation block is a hallmark of AML. However, whether an analogous block exists in CMML monocyte and granulocyte development is unknown. Although alterations in monocyte activation are reported in CMML, whether CMML monocytes are defective in differentiation to macrophages *in vivo* is unclear^6^. Recently, we have shown that CMML monocytes could be successfully differentiated to macrophages *in vitro*^10^. This suggests that any monocyte maturation defects in CMML are likely cell extrinsic. Treating healthy monocytes with resistin prevented monocyte differentiation to macrophages, presumably by downregulating key activation genes such as *CD163*. Whether elevated levels of plasma resistin indeed affects monocyte maturation in CMML patients *in vivo*, and thereby contributes to the expansion of monocytes, remains unknown and difficult to empirically test. We and others have shown that Defensin levels are elevated in CMML granulocytes at both RNA and protein level^6^. Granulocyte derived defensins have been shown to affect monocyte maturation, and here we show that resistin also induces expression of defensins in monocytes. Presumably, Resistin could also induce the expression of defensins in neutrophils in an autocrine manner, potentially further contributing to defects in monocyte maturation.

We identified elevated frequencies of immune suppressive cell types in most CMML patients tested. Elevated levels of Tregs are common in many myeloid cancers including CML, AML, MPN and MDS and are associated with worse prognosis^52^. In MDS, higher levels of Tregs and MDSCs are linked to disease progression and increased risk of AML transformation^53,54^. This is further supported by the observation that immune suppression following solid organ transplantation increases the risk of leukemic development, suggesting a causal link between immune suppression and leukaemia onset^55^. A positive correlation between MDSC and Treg levels in myeloid cancers suggest a crosstalk between these two populations. In this study, we showed that higher plasma resistin levels are associated with risk of AML transformation. It is tempting to speculate that resistin produced by MDSCs contributes to changes in T cells profiles, thereby promoting leukaemia development.

We propose that the immune suppression observed in CMML patients is, in part, mediated by SEMA4A. We observed that CMML monocyte exhibit elevated levels of *SEMA4A*, and its expression is induced by resistin. SEMA4A has been shown to bind to NRP1 (in mice) or PlexinB1(in humans) on Tregs, and stabilizes T-reg cell populations, thereby inducing immune tolerance against tumours^48,56^. In addition, human SEMA4A has also been shown to bind to ILT4 on CD4+ T cells and promote Th2 responses^49^. Indeed, SEMA4A knockout mice show defects in Th1/Th2 regulation^57^. Targeting SEMA4A using antibody drug conjugates could potentially alleviate immune suppression and improve patient outcomes^58^.

In humans, elevated plasma resistin levels are an independent predictor of atherosclerosis, and resistin is known to aggravate cardiovascular events through multiple mechanisms^59^. Notably, a significant percentage of CMML patients (∼36%) experience cardiovascular complications^60,61^. Future research investigating the relationship between plasma resistin levels with cardiovascular events in CMML patients could have value for patient monitoring and preventative management. Immature granulocytes are also frequently observed in solid tumors^62^. The frequency of LDNs, a population comprising immature MDSCs and mature neutrophils, in the PB of solid cancer patients, is associated with poor prognosis^63,64^. Furthermore, TANs often exhibit immature and immune suppressive phenotypes that promote tumour growth^65^. Investigating whether TANs release resistin and contribute to immune suppressive features in solid cancers patients is a future area of interest. Therapeutic strategies aimed at promoting neutrophil maturation, shifting their phenotype from N2 to N1 type, or depleting immature granulocytes altogether may offer potential benefits for cancer patients with elevated LDNs. For instance, FATP2, a key driver of MDSC phenotype, has demonstrated therapeutic promise^66^. Alternately targeting Syk could potentially eliminate neutrophils. Notably, clinical trials using Fostamatinib, a Syk inhibitor, are currently underway for CMML patients.

## Materials and Methods

### Patients and Samples

CMML samples were obtained from patients treated at the Christie NHS Foundation Trust (Manchester UK) and recruited to the Manchester Cancer Research Centre Haematological Malignancies Tissue Biobank. Healthy volunteers were used as controls. All patients and volunteers gave informed consent according to the Declaration of Helsinki. The study was approved by the UK Health Research Authority (REC: 19/LO/0564).

### Granulocyte Isolation and Characterisation

Granulocytes were isolated from 2 – 5 ml PB of healthy and CMML donors using EasySep™ Direct Human Neutrophil Isolation Kit (Stem Cell Technologies, 19666). Isolated granulocytes were washed twice in SM buffer (phenol red-free RPMI (Sigma-Aldrich, R7509) containing 3% heat inactivated (HI) FBS (Gibco, A5256701) and 4 mM EDTA (Invitrogen, 15575-038) and counted. From this, 1 x 10^5^ cells were stained with CD14-FITC (Invitrogen, 11-0149-42), CD16-APC-H7 (BD Biosciences, 560195), CD13-PE-Cy7 (BD Biosciences, 561599), CD11b-PE (Invitrogen, 12-0118-42) and CD66b-APC (Invitrogen, 17-0666-42) to profile purity and maturation status of granulocytes. Cells were analysed on a BD LSR II flow cytometer. In total, we have analysed granulocytes from 19 healthy volunteers and 26 CMML patients in this study. Flow data (from 12 controls and 12 *TET2* mutant CMML patients) from our previous study^16^ was reanalysed and used in this study.

### Granulocyte RNA extraction and RNA

RNA was extracted from 5 x 10^5^ granulocytes from 14 healthy volunteers (7 young, 7 old) and 14 CMML donors using the RNeasy Micro Plus kit (QIAGEN, 74034), according to manufacturer’s recommendations.

### Phagocytosis assay

A total of 1 x 10^5^ granulocytes from three aged healthy and six CMML donors were resuspended in HBSS (Gibco, 14025-092) containing 10 mM HEPES (Gibco, 15630-080) and 3% HI FBS and incubated at 37°C for 30 minutes with/without Zymosan A (S. cerevisiae) BioParticles, Alexa Fluor 488 conjugated, (ThermoFisher, Z23373) at MOI 10:1, according to manufacturer’s protocol. Activity was stopped by adding ice-cold PBS, centrifuging 300 x g for 5 mins at 4°C and washing twice in ice-cold PBS. Bioparticle uptake was measured by flow cytometry on BD FACS Canto instrument.

### Resistin ELISA

Platelet poor plasma (PPP) was isolated from twenty-seven CMML and fourteen healthy donor PB by centrifuging blood at 200 x g for 10 minutes. The plasma supernatant was removed and added to 1 ml centrifuge tube and centrifuged at 2000 x g for 10 minutes. The supernatant was added to a fresh tube and centrifuged a final time at 2000 x g for 10 minutes, leaving a PPP plasma supernatant. The PPP supernatant was stored at -80°C. The resistin concentration was measured in the PPP supernatant using the Human Resistin Standard ABTS ELISA Development Kit (Peprotech, 900-K235), according to manufacturer’s protocol. Absorbance was measured at 440 nm using BMG FLUOstar Omega plate reader and resistin concentration calculated using a standard curve created on GraphPad Prism software.

### Monocyte isolation and resistin treatment

PB from aged healthy volunteers was diluted 1:1 with PBS and underwent density gradient separation by layering onto an equal volume of Ficoll-Paque Plus (Cytiva, 17144003) and centrifuged at 400 x g for 20 minutes at 20°C with no brake deceleration and acceleration set at 3 in Eppendorf 5810R centrifuge. The white layer containing mononuclear cells (MNCs) was collected and incubated for 3 minutes at RT in ACK-lysis buffer (Gibco, A1049201) for red blood cell removal. Cells were washed twice in PBS and counted. CD14^+^ monocytes were isolated by magnetic activated cell sorting (MACS) using CD14 microbeads (Miltenyi, 130-050-201), following manufacturer’s protocol. Isolated CD14^+^ monocytes were seeded out at 1 x 10^5^ cells/ml in 24 well plates in monocyte complete medium - RPMI media (Gibco, 21875) containing 10% HI FBS, 1% penicillin-streptomycin (Sigma-Aldrich, P0781), and 10 ng/ml M-CSF (Peprotech, 300-25) - with GFP control (Abcam Ltd, ab84191) or resistin (Cambridge Bioscience, 230-00271-100) and incubated at 37°C, 5% CO_2_ for 24- and 48-hours. Following incubation, cells were harvested and stained with CD14-FITC and CD16-APC-H7 or SEMA4A-APC (BioLegend, 148405) and analysed by flow cytometry on BD FACS Canto.

### Macrophage differentiation

CD14^+^ monocytes were isolated from four healthy volunteers and seeded out at 1 x 10^6^ cells/ml in a 6 well plate in monocyte complete medium containing 25 ng/ml M-CSF and exposed to either 250 ng/ml GFP endotoxin control or recombinant resistin in duplicate. Cells were incubated for 6 days at 5% CO_2_ and 37°C, with fresh complete medium containing treatment added on Day 3. On Day 6 cell confluency of attached cells was calculated using the AI Cell Confluency module (Labscope software) of a Zeiss Axiovert 5 digital microscope. Media was harvested and the number of viable suspended cells calculated using trypan blue and an Invitrogen Countess 3 Cell Counter.

### Monocyte TLR4 treatment

CD14^+^ monocytes from four aged healthy volunteers were seeded out at 2 x 10^5^ cells/ml in 48 well plates with monocyte complete medium and pretreated for 1 hour with/without 10 μg/ml TLR4 neutralising antibody (R&D systems, AF1478) at 37°C, 5% CO_2_. After pretreatment, cells were treated with either GFP control or 250 ng/ml resistin and incubated for 24- and 48-hours. Following incubation cells were stained with CD14-FITC and CD16-APC-H7 antibodies and analysed by flow cytometry on BD FACS Canto.

### Monocyte RNA extraction and RNA sequencing

CD14^+^ monocytes from four aged healthy volunteers were seeded at 5 x 10^5^ cells/ml in 12-well plates with monocyte complete medium and treated with either GFP, 250 ng/ml resistin or 100 ng/ml LPS (Sigma-Aldrich L2630) for 6 or 24 hours. Cells were then harvested, washed in PBS and RNA extracted using RNeasy Micro Plus kit. RNA was then sequenced 3’ single end using the Novaseq SP instrument 1 x 100 cycles.

### Apoptosis assays

Six cryopreserved CMML PBMNCs were thawed and non-viable cells removed by dead cell removal kit (Miltenyi, 130090101), following manufacturer’s protocol. CD14+ cells were isolated by MACS and cultured in monocyte complete medium containing either GFP or 250 ng/ml resistin for 72hrs at a density of 1 x 10^5^ cells/ml in a 24-well plate. Following incubation, cells were harvested and washed once in PBS and then once in 1x annexin V binding buffer (AVBB) (Thermo Fisher Scientific, V13246). Cells were resuspended in AVBB containing Annexin V-APC (BioLegend, 640941) and incubated at RT for 15 minutes, protected from light. The cell suspension was then washed with AVBB and centrifuged at 400 x g for 5 minutes. Cells were resuspended in AVBB containing 7-AAD viability stain (BioLegend, 420404) and incubated at RT for 15 minutes. Following incubation, the cell suspension was topped up with AVBB and analysed by flow cytometry on BD Fortessa.

### CellTiter glo assays

Eight cryopreserved CMML PBMNCs were thawed and non-viable cells removed by dead cell removal kit. CD14+ cells were isolated by MACS and cultured in monocyte complete medium containing either GFP or 10, 25, 100 or 250 ng/ml resistin for 72hrs at a density of 1 x 10^4^ cells/ml in a 96-well plate. Following incubation, equal volumes of CellTiter-Glo (Promega, G9243) was added to the cell cultures, following manufacturer’s recommendations. The relative luminescence was then measured using BMG Labtech FLUOstar Omega plate reader.

### Treg populations in healthy and CMML PBMNCs

Fresh PBMNCs were isolated from eight healthy volunteers using density gradient separation and CD14+ monocytes isolated by MACS, as previously described, and CD14^-^ cells were used for measuring the percentage of Tregs. Separately, thirteen cryopreserved CMML patients’ PBMNCs were thawed. A total of 1 x 10^6^ CMML PBMNCs and CD14^-^ healthy PBMNCs were stained with CD45-BV785 (BioLegend, 304048), CD3-PerCP-CY5.5 (BioLegend, 300430), CD56-APC-Cy7 (BioLegend, 318332), CD8a-AF700 (BioLegend, 300920), CD4-BV605 (BioLegend, 317438), CD127-PE-Cy7 (BD biosciences, 560822), CD25-APC (BioLegend, 302610) and FOXP3-PE (Invitogen, 12-4776-42). Cells were analysed by flow cytometry on BD Fortessa and percentage of Treg cells was determined using FlowJo software.

### PBMC co-culture with resistin

PBMNCs with or without depletion of CD14^+^ cells were seeded at 1 x 10^6^ cells/ml in 6 well plates with T-cell complete medium - RPMI supplemented with 10% HI FBS, 1% penicillin-streptomycin, 25 ng/ml M-CSF, 20 IU/ml IL2 (Chiron), 10 ng/ml IL7 (Peprotech, 200-07), 1 mM sodium pyruvate (Corning, 25-000-CI), 1% MEM non-essential amino acid solution (Sigma-Aldrich, M7145), 25 mM HEPES and 50 nM 2-mercaptoethanol (Sigma-Aldrich, M6250) - and treated with GFP control or 250 ng/ml resistin for 9 days. Fresh T-cell complete medium containing GFP or resistin was added every 3 days. After the incubation period, cells were harvested, washed in SM buffer, counted and 1 x 10^6^ cells were stained with Zombie UV viability stain (BioLegend, 423108), CD3-PerCP-CY5.5, CD4-BV605, CD8a-AF700; for Tregs the additional stains CD45-BV785, CD56-APC-Cy7, CD127-PE-CY7, CD25-APC and FOXP3-PE were used; for Th1/Th2 the additional stains CD183-BV421 (BioLegend, 353715), CD194-PE-Cy7 (BioLegend, 359409), CD196-PE (BioLegend, 353409) and CCR10-APC (BioLegend, 341505) were used. Cells were analysed by flow cytometry on BD Fortessa and percentage of Treg or Th1 and Th2 cells was determined using FlowJo software.

### Th2/Th1 ratio in CMML samples

Eight cryopreserved CMML PBMNCs were thawed and non-viable cells removed by dead cell removal kit. CD14^+^ cells were removed by MACS and 1 x 10^6^ CD14^-^PBMC cells were stained with live/dead-near IR (ThermoFisher, L34975), CD3-AF700 (BioLegend, 300323), CD4-FITC (BioLegend, 300505), CD8a-BV510 (BioLegend, 301047), CD183-BV421, CD194-PE-CY7, CD196-PE and CCR10-APC. Cells were analysed by flow cytometry on BD Fortessa and percentage of Th1 and Th2 cells was determined using FlowJo software.

### Transcriptome analysis (Granulocyte data)

Sample libraries were prepared by 3’ Quantseq (Lexogen) protocol and were sequenced using the Novaseq (Illumina) platform. Basecalls were converted to fastq files using bcl2fastq (Illumina). Fastq files were aligned to GRCh38 using Star aligner (version 2.5.1b; Ensembl85). BAM alignments were quantified in R (version 3.6.1) using featureCounts from the Rsubread library (version 2.0.1) and the Ensemble GTF annotation (Homo_sapiens.GRCh38.85.gtf). Differential expression (DE) was evaluated comparing the gene level integer read count data for between patient/sample groups using the DESeq2.v 1.26.0 Bioconductor package. DESeq2 DEG estimation involves multiple steps. In brief, it uses normalization factor to model read counts to account for sequencing depth. Then it estimates gene-wise dispersions and perform shrinkage analysis to generate more accurate estimates of dispersion to model the counts. Finally, it fits the negative binomial model and performs hypothesis testing using the Wald test.

### Transcriptome analysis (Monocyte data)

Quality of FASTQ files was evaluated using the FASTQC (v.0.12.1) and MULTIQC (v.1.15) software. MULTIQC adapter content check passed with no warnings, suggesting against need for adapter trimming. Reads were mapped to the reference GENCODE transcriptome (release v42; GRCh38.p13/hg38) using the STAR (v.2.5.1b) aligner with the parameter *–quantMode TranscriptomeSAM*. The resulting *Aligned.toTranscriptome.out bam* files from STAR were then used as impute for transcripts counts quantification via Salmon (v.1.10.2) using the alignment-based mode and the following parameters: *--libType A, --gcBias, --seqBias*. The following downstream analysis was then performed using R (v.4.3.3). Gene-level abundances (TPMs) and counts were estimated via using the package TxImport (v.1.30.0). Transcript-level counts contained within Salmon-produced *quant.sf* files were processed with the *tximport* function, setting parameter *countsFromAbundance =”no”* for subsequent DESeq2 (v.1.42.1) offset method and using a reference transcript to gene dictionary. This was generated from a txDb object obtained via GenomicFeatures (v.1.54.4) package using the function *makeTxDbFromGFF* on a GENCODEv42 gtf reference file.

Differentially expressed genes (DEGs) were computed using DESeq2 importing gene counts with the *DESeqDataSetFromTximport* function. Following library size estimation, genes with <5 counts across a minimum of 3 samples were removed. A minority (0.002%) of Ensembl genes mapping to the same gene symbol were also filtered to maintain only the most informative duplicate with the highest base mean expression. Following filtering, *DESeq2* function was run using *’local’* fit to model gene expression, since this resulted in the best fitting model with the least residuals. Finally, counts were normalized by condition (*design =* ∼*Condition*) and transformed using the regularized log transformation (rlog), which accounts for gene dispersion and results in a log2 scaled count matrix. DEG analysis was performed implementing the independent hypothesis weighting (IHW) pvalue correction to increase detection rate. Results were sorted by increasing p.adjusted value. Rlog counts were used for visualisation purposes, including heatmap representations.

Gene set enrichment analysis (GSEA) (v.4.3.3) was used^67,68^. The normalized expression values was used as an input for this analysis. Gene set permutation as opposed to phenotype permutation was used, due to sample size, to calculate significance. 1000 permutations were used.

### Survival analysis

Survival analyses were performed with the Kaplan–Meier method, and differences between survival curves were assessed using the log-rank test. AML-free survival (LFS) was defined as the interval from initial diagnosis to AML transformation, death, or the last follow-up, while overall survival (OS) was defined as the time from diagnosis to the last follow-up or death from any cause. All statistical analyses and data visualisations were carried out using GraphPad Prism (version 10.0.2) and R software (version 4.3.1).

### Statistical analysis

Categorical and nominal variables were analysed using either the chi-squared test or Fisher’s exact test, as appropriate. Continuous variables were compared using the Mann– Whitney test or the Kruskal–Wallis test, as appropriate. The optimal cut-off for distinguishing patient groups was determined using maximally selected rank statistics from the maxstat R package.

Data availability: Sequencing data were deposited into the Gene Expression Omnibus database under accession number **GSE289009**

## Acknowledgements

The Epigenetics of Haematopoiesis group is funded by The Oglesby Charitable Trust. We thank Cancer Research UK Manchester Institute core facilities including molecular biology core facility and flow facility. We thank all the members of Epigenetics of Haematopoiesis group for helpful discussion and technical support. All illustrations in the manuscript were created with BioRender.com.

## Author contributions

K.B., D.W. & N.J.H conceived and designed the study. K.B., N.J.H., R.C., L.A.G., K.G., Y.H.W., & H.H.E. performed all the experiments and analysed the data. E.S., D.S., D.B., C.L., H.T., provided advice on experimental design, access to datasets, and critical feedback on the manuscript. K.B., D.W. & N.J.H wrote the manuscript.

