## Supplementary table 1 for "Granulocyte Derived Resistin Inhibits Monocyte Maturation and Induces Immune Suppression in CMML"

| Healthy ID | Age (Years) | Sex | Included in |
| --- | --- | --- | --- |
| YHB1 | 30 | F | Granulocyte flow, Granulocyte RNA-seq |
| YHB2 | 30 | F | Granulocyte flow, Granulocyte RNA-seq |
| YHB3 | 24 | F | Granulocyte flow, Granulocyte RNA-seq |
| YHB4 | 21 | M | Granulocyte flow, Granulocyte RNA-seq |
| YHB5 | 33 | M | Granulocyte flow, Granulocyte RNA-seq, Treg co-culture, Th1/Th2 co-culture, Th2/Th1 ratio, Treg |
| YHB6 | 26 | M | Granulocyte flow, Granulocyte RNA-seq |
| YHB7 | 24 | M | Granulocyte flow, Granulocyte RNA-seq, Treg co-culture, Th1/Th2 co-culture, Th2/Th1 ratio, Treg |
| HB1 | 49 | F | Granulocyte flow, Granulocyte RNA-seq |
| HB6 | 51 | F | Plasma Resistin Elisa |
| HB9 | 71 | F | Granulocyte flow, Granulocyte RNA-seq, Plasma Resistin Elisa |
| HB10 | 58 | F | Granulocyte RNA-seq, Plasma Resistin Elisa |
| HB11 | 64 | F | Granulocyte flow, Granulocyte RNA-seq, Plasma Resistin Elisa |
| HB12 | 63 | F | Granulocyte flow, Granulocyte RNA-seq, Plasma Resistin Elisa |
| HB13 | 52 | M | Granulocyte flow, Granulocyte RNA-seq, Plasma Resistin Elisa |
| HB14 | 55 | F | Granulocyte RNA-seq, Plasma Resistin Elisa |
| HB17 | 55 | M | Plasma Resistin Elisa |
| HB18 | 21 | F | Plasma Resistin Elisa |
| HB19 | 55 | F | Granulocyte flow, Plasma Resistin Elisa |
| HB20 | 56 | M | Plasma Resistin Elisa |
| HB21 | 58 | M | Plasma Resistin Elisa |
| HB22 | 60 | M | Plasma Resistin Elisa |
| HB23 | 53 | F | Plasma Resistin Elisa |
| HB24 | 60 | M | Monocyte classical repartitioning |
| HB28 | 53 | M | Granulocyte flow, TLR4 nAb, Monocyte RNA-seq |
| HB29 | 58 | F | Granulocyte flow, TLR4 nAb, Monocyte RNA-seq |
| HB30 | 33 | M | TLR4 nAb, Monocyte RNA-seq |
| HB31 | 58 | M | Granulocyte flow, TLR4 nAb, Monocyte RNA-seq |
| HB32 | 39 | F | Monocyte differentiation, Th2/Th1 ratio, Treg |
| HB33 | 28 | M | Monocyte differentiation, Th2/Th1 ratio, Treg |
| HB34 | 40 | F | Monocyte differentiation, Th2/Th1 ratio, Treg |
| HB35 | 47 | F | Monocyte differentiation, Treg co-culture, Th1/Th2 co-culture, Th2/Th1 ratio, Treg |
| HB36 | 54 | M | Granulocyte flow, Phagocytosis, Monocyte classical repartitioning, resistin exposed monocytes SEMA4A expression |
| HB37 | 65 | M | Granulocyte flow, Phagocytosis, Monocyte classical repartitioning, resistin exposed monocytes SEMA4A expression |
| HB38 | 51 | M | Granulocyte flow, Phagocytosis, Monocyte classical repartitioning, resistin exposed monocytes SEMA4A expression |
| HB39 | 38 | M | Th2/Th1 ratio, Treg |
| HB40 | 57 | F | Monocyte classical repartitioning, resistin exposed monocytes SEMA4A expression |
| HB41 | 24 | F | Treg co-culture, Th1/Th2 co-culture, Th2/Th1 ratio, Treg |
| HB49 | 41 | F | Treg co-culture, Th1/Th2 co-culture, Th2/Th1 ratio |
| HB50 | 25 | F | Treg co-culture, Th1/Th2 co-culture, Th2/Th1 ratio |

| Patient ID | Age (Years) | Sex | Type | Stage | Mutations | Included in |
| --- | --- | --- | --- | --- | --- | --- |
| 547 | 78.4 | M | Pro | 2 | SRSF2, TET2, ASXL1, RUNX1 | Plasma Resistin Elisa, Apoptosis, Cell viability, Th2/Th1 ratio, Treg |
| 572 | 60.0 | M | Pro | 2 | TET2, ASXL1, ZRSR2, RUNX1, KRAS, BCOR | Plasma Resistin Elisa |
| 612 | 66.5 | M | Dys | 1 | SRSF2, TET2, KRAS | Granulocyte flow, Granulocyte RNA-seq, Plasma Resistin Elisa, Resisn vs *RETN* correlation, Resistin vs flow parameters |
| 620 | 67.8 | M | Dys | 2 | SF3B1, DNMT3A | Plasma Resistin Elisa, Apoptosis, Cell viability, Th2/Th1 ratio, Treg |
| 626 | 83.6 | M | Dys | 2 | TET2, U2AF1, DNMT3A, RUNX1, KRAS, BCOR | Plasma Resistin Elisa, Apoptosis, Cell viability, Th2/Th1 ratio, Treg |
| 628 | 59.7 | F | Dys | 1 | SRSF2, TP53 | Plasma Resistin Elisa |
| 666 | 63.4 | F | Dys | 1 | SRSF2, TET2, ASXL1, CBL | Plasma Resistin Elisa, Apoptosis, Cell viability, Th2/Th1 ratio, Treg |
| 786 | 74.2 | M | Pro | 1 | SRSF2, TET2, ASXL1, CBL | Plasma Resistin Elisa, Th2/Th1 ratio, Treg |
| 830 | 65.3 | M | Dys | 1 | TET2, NRAS, CBL | Granulocyte flow, Granulocyte RNA-seq, Plasma Resistin Elisa, Resisn vs *RETN* correlation, Resistin vs flow parameters, Treg |
| 909 | 58.2 | M | Pro | 1 | ASXL1, KIT | Granulocyte flow, Plasma Resistin Elisa, Resistin vs flow parameters |
| 986 | 66.2 | F | Dys | 1 | TET2, KRAS, NRAS, TP53, CBL | Granulocyte flow, Granulocyte RNA-seq |
| 999 | 63.1 | M | Pro | 1 | TET2, U2AF1, PHF6, CBL | Granulocyte flow, Granulocyte RNA-seq |
| 1092 | 61.4 | M | Dys | 1 | ASXL1, KRAS | Granulocyte flow |
| 1107 | 75.3 | F | Pro | 1 | JAK2, TET2, ASXL1, CBL | Plasma Resistin Elisa |
| 1115 | 77.2 | M | Pro | 1 | SRSF2, TET2, RUNX1, NRAS | Granulocyte flow, Granulocyte RNA-seq, Plasma Resistin Elisa, Resisn vs *RETN* correlation, Resistin vs flow parameters, Apoptosis, Cell viability, Th2/Th1 ratio, Treg |
| 1122 | 61.5 | M | Pro | 1 | SRSF2, TET2, ASXL1, RUNX1, NRAS | Granulocyte flow, Phagocytosis, Plasma Resistin Elisa, Resistin vs flow parameters, Treg |
| 1172 | 67.0 | M | Dys | 1 | SRSF2, TET2 | Granulocyte flow, Granulocyte RNA-seq |
| 1188 | 80.5 | F | Pro | 1 | SRSF2, TET2, KRAS | Granulocyte flow, Granulocyte RNA-seq, Phagocytosis, Plasma Resistin Elisa, Resisn vs *RETN* correlation, Resistin vs flow parameters, Treg |
| 1206 | 71.2 | M | Pro | 1 | SF3B1, RUNX1 | Granulocyte flow, Granulocyte RNA-seq |
| 1213 | 53.8 | M | Pro | 1 | SRSF2, TET2, KRAS, NRAS | Granulocyte flow, Phagocytosis, Plasma Resistin Elisa, Resistin vs flow parameters, Cell viability |
| 1214 | 75.2 | M | Dys | 1 | SRSF2, IDH2, KRAS | Granulocyte flow, Granulocyte RNA-seq |
| 1216 | 65.4 | M | Dys | 1 | TET2 | Granulocyte flow, Granulocyte RNA-seq |
| 1223 | 73.0 | F | Pro | 1 | SRSF2, TET2, EZH2 | Granulocyte flow, Granulocyte RNA-seq, Plasma Resistin Elisa, Resisn vs *RETN* correlation, Resistin vs flow parameters, Th2/Th1 ratio, Treg |
| 1236 | 64.3 | M | Pro | 1 | ASXL1, EZH2, FLT3 | Granulocyte flow, Plasma Resistin Elisa, Resistin vs flow parameters |
| 1238 | 74.5 | M | Pro | 1 | SRSF2, TET2, ASXL1, RUNX1, KIT | Granulocyte flow, Granulocyte RNA-seq, Plasma Resistin Elisa, Resisn vs *RETN* correlation, Resistin vs flow parameters, Treg |
| 1239 | 69.8 | F | Pro | 1 | TET2, ASXL1, NRAS | Granulocyte flow, Granulocyte RNA-seq, Plasma Resistin Elisa, Resisn vs *RETN* correlation, Resistin vs flow parameters, Apoptosis, Cell viability, Th2/Th1 ratio, Treg |
| 1241 | 63.4 | M | Pro | 1 | JAK2, TET2, ASXL1, NPM1 | Plasma Resistin Elisa, Treg |
| 1242 | 75.5 | F | Pro | 1 | TET2, NRAS | Granulocyte flow, Granulocyte RNA-seq, Plasma Resistin Elisa, Resisn vs *RETN* correlation, Resistin vs flow parameters |
| 1243 | 71.6 | M | Pro | 1 | ASXL1, SETBP1 | Granulocyte flow, Plasma Resistin Elisa, Resistin vs flow parameters |
| 1244 | 59.4 | M | Dys | 1 | SRSF2, TET2, CBL | Granulocyte flow, Phagocytosis, Plasma Resistin Elisa, Resistin vs flow parameters |
| 1255 | 75.8 | F | Pro | 1 | TET2, CBL | Granulocyte flow, Plasma Resistin Elisa, Resistin vs flow parameters |
| 1266 | 72.5 | M | Pro | 1 | SRSF2, TET2, CBL | Granulocyte flow, Plasma Resistin Elisa, Resistin vs flow parameters |
| 1269 | 84.7 | M | Pro | 1 | SRSF2, ASXL1, IDH2 | Granulocyte flow, Plasma Resistin Elisa, Resistin vs flow parameters |
| 1271 | 72.8 | F | Dys | 1 | TET2, ASXL1, NRAS, PHF6 | Granulocyte flow, Phagocytosis, Plasma Resistin Elisa, Resistin vs flow parameters |
| 1290 | 73.2 | M | Pro | 1 | ASXL1, DNMT3A, RUNX1, KRAS | Granulocyte flow, Phagocytosis, Resistin vs flow parameters, Cell viability |
