## Supplementary data for "Granulocyte Derived Resistin Inhibits Monocyte Maturation and Induces Immune Suppression in CMML"

**Supplementary tables:**

**Table S1:** Patient information

**Table S2:** Genes upregulated or down regulated in CMML granulocytes compared to all granulocytes

**Table S3:** Genes upregulated or down regulated in old granulocytes compared to young granulocytes.

**Table S4:** Genes upregulated or down regulated in healthy monocytes treated with resistin for 6 hrs in comparison with GFP control.

**Table S5:** Genes upregulated or down regulated in healthy monocytes treated with resistin for 24 hrs in comparison with GFP control.

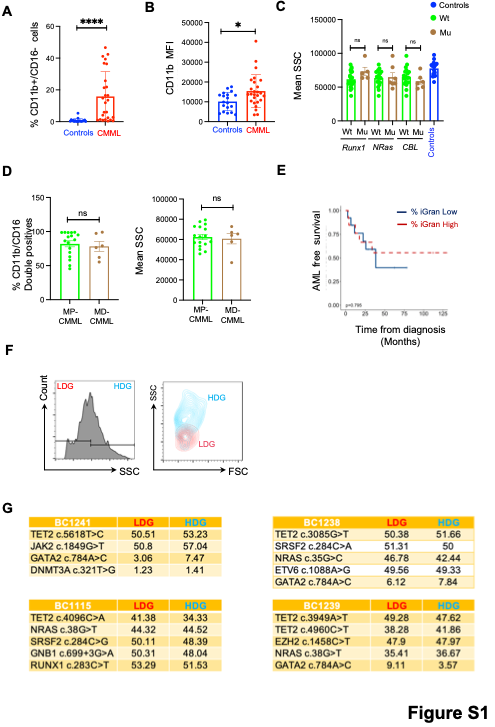

**Figure S1: A.** Bar graph showing average percentage of CD11b^+^/CD16^-^ cells in control (N=19) and CMML (N=26) granulocytes. **B.** Average mean fluorescence intensity (MFI) of cell surface marker CD11b in healthy (N=19) and CMML granulocytes (N=26). **C.** Bar graph showing average granulocyte SSC in wild-type and mutant genotype of indicated genes recurrently observed in CMML patients. **D**. Bar graph illustrating percentage of CD11b/CD16 double positive mature granulocytes or mean granulocyte SSC-A between CMML disease subtypes dysplastic (MD-CMML) and proliferative (MP-CMML). **E.** Kaplan-Meier AML-free survival plot of CMML patients stratified by the percent of immature granulocytes (iGran). High = > 13% (N=13) vs low = <13% (N=13). **F.** A representative flow cytometry plot (left) displaying gating strategy used to sort low-density granulocytes (LDGs) and high-density granulocytes (HDGs). A representative flow cytometry plot (right) displaying the side scatter profile of LDGs and HDGs sorted from CMML patient granulocytes (N=4). **G**. Tables displaying the variant allele frequencies of indicated mutations in LDGs and HDGs sorted from 4 different CMML patient granulocytes. Error bars = ± SEM; ns, no significance; *, P<0.05; **, P<0.01; ****, P<0.0001.

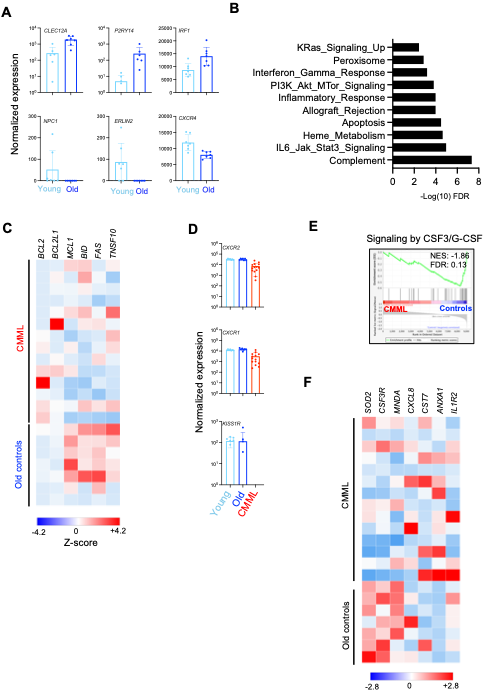

**Figure S2: A.** Normalized expression levels of indicated genes in young and old healthy granulocytes. **B.** GO analysis of downregulated genes in CMML granulocytes as compared with controls. Top ten significant pathways are shown. **C.** Heatmap showing expression levels of indicated pro- or anti-apoptotic genes in old control and CMML granulocytes. **D.** Normalized expression levels of selected downregulated genes in CMML granulocytes as compared with healthy granulocytes. **E.** GSEA plots showing relative enrichment of a gene set linked to CSF3/G-CSF signalling. **F.** Heatmap showing expression levels of indicated genes related to neutrophil activation in old control and CMML granulocytes.

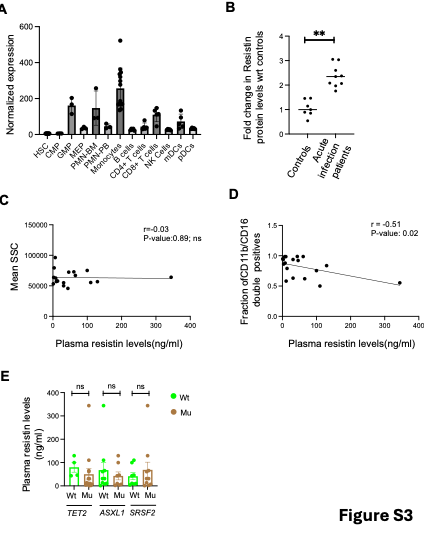

**Figure S3:** **A.** Relative expression levels of *RETN* in different normal haematopoietic cell types. Data plotted from Bloodspot. **B.** Fold change in plasma resistin levels in individuals experiencing severe infections as compared with healthy controls^26^. **C & D.** Pearson correlation between plasma resistin levels with granularity (C) or fraction of CD11b/CD16 double positive population (D) in CMML patients. **E.** Bar graph showing average plasma resistin levels in wild-type and mutant genotype of indicated recurrently mutated genes seen in CMML patients. Error bars = ± SEM; **, P<0.01.

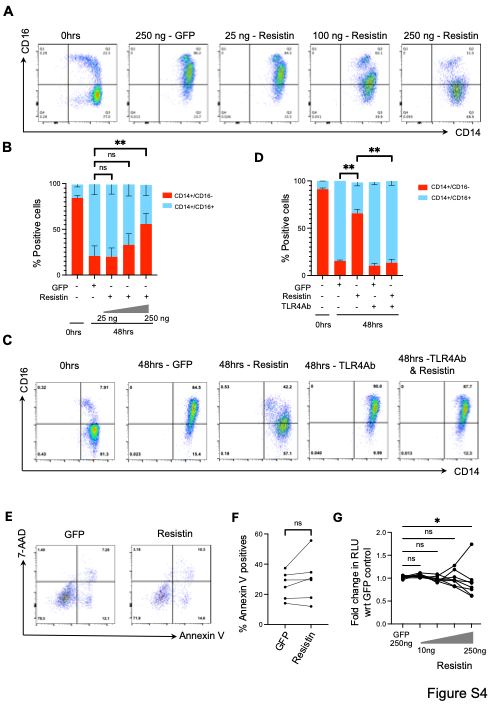

**Figure S4: A.** Representative flow cytometry plots showing the surface marker expression of CD14 and CD16 on freshly isolated healthy monocytes (0hrs) and monocytes treated with increase in concentration of resistin or control GFP for 48 hours (N=5). **B & D.** Bar charts showing average percentage of CD14+ or CD14+CD16+ populations under indicated conditions. **C.** Representative flow cytometry plots showing the surface marker expression of CD14 and CD16 on monocytes incubated with or without TLR4 neutralising antibody (TLR4Ab) prior to exposure with resistin or GFP control for 48 hours (N=4). **E.** Representative flow cytometry plots displaying Annexin V/7AAD levels on CMML monocytes treated with resistin or GFP for 72 hours. **F.** Dot plot showing the percentage of Annexin V positive cells when CMML monocytes are treated with resistin in comparison with GFP control (N=6). **G.** Cell viability determined by CellTiter-Glo assay of CMML monocytes treated with either increase in concentration of resistin (10 ng to 250 ng) or control GFP (250 ng/ml) for 72 hours (N=8). Fold change in relative luminescence units (RLU) in monocytes treated with resistin with respect to GFP control is shown. Error bars = ± SEM; ns, no significance; *, P<0.05; **, P<0.01.

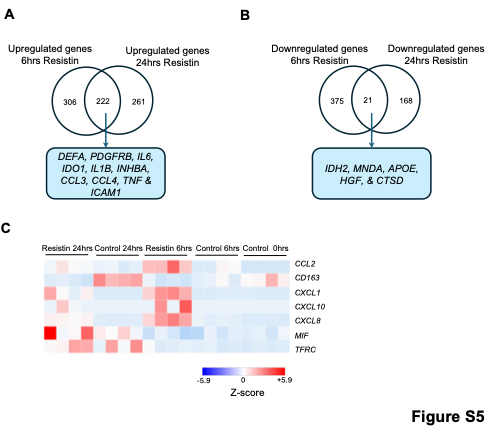

**Figure S5: A.** Venn diagram showing overlap of differentially expressed genes upregulated following 6 hrs and 24 hrs treatment with resistin in comparison with their respective time point controls. **B.** Venn diagram showing overlap of differentially expressed genes downregulated following 6 hrs and 24 hrs treatment with resistin in comparison with their respective time point controls. **C.** Heatmap showing expression of different genes related to monocyte activation in different experimental conditions as indicated.

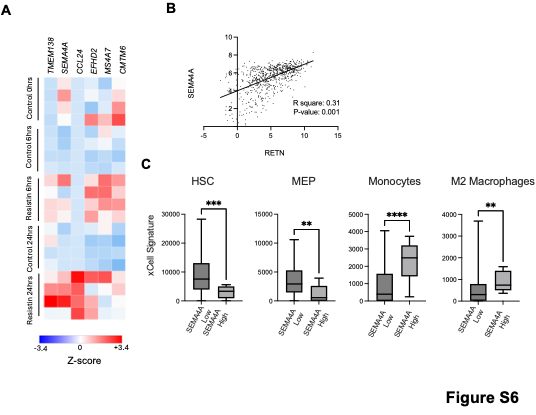

**Figure S6: A.** Heatmap showing relative expression of indicated genes commonly upregulated at 6 and 24 hrs of resistin treatment but not by LPS. **B.** Correlation between expression levels of *RETN* and *SEMA4A* in AML samples^47^. **C.** Cybersort analysis showing relative enrichment of indicate haematopoietic cell signatures in *SEMA4A* high vs low CMML patients (N=11 High; N=79 Low). HSC: Haematopoietic stem cells. MEP: Megakaryocyte erythroid progenitors. Error bars = ± SEM; **, P<0.01; ***, P<0.001; ****, P<0.0001

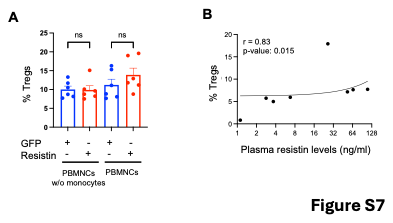

**Figure S7:** **A.** Percentage of Treg cells in ex vivo cultures of whole- or CD14 depleted-PBMNCs treated with either GFP or resistin for 9 days (N=6). **B.** Correlation plot of resistin plasma concentration against percentage of Tregs in CMML patients’ PB.
